# Stress-induced plasminogen activator inhibitor-1 (PAI-1) as a blood biomarker and brain risk factor for PTSD

**DOI:** 10.1101/2025.07.02.661922

**Authors:** M Mennesson, S Abdelkaoui, V Roullot-Lacarriere, S Tronel, A Cathala, V Lalanne, PL Raux, L Makrini, E Valjent, AM Duffaud, D Claverie, M Vallée, A Desmedt, M Trousselard, JM Revest

**Affiliations:** Univ. Bordeaux, Inserm, Neurocentre Magendie, U1215, F-33000 Bordeaux, France; IGF, Université de Montpellier CNRS, Inserm, 34094 Montpellier, France; INM, Univ. Montpellier, CNRS, Inserm, 34094 Montpellier, France; Unité de Neurophysiologie du Stress, Département Neurosciences & Contraintes Opérationnelles, Institut de Recherche Biomédicale des Armées (IRBA), 91223 Brétigny-sur-Orge, France; Réseau ABC des Psychotraumas, Montpellier, France; École de Psychologues Praticiens (EPP), 23 rue du Montparnasse, 75006 Paris, France; Université de Lorraine, Inserm, INSPIIRE, UMR 1319, F-54000, Nancy

## Abstract

Post-traumatic stress disorder (PTSD) is a severe stress-related psychiatric condition triggered by traumatic life-threatening events, characterized notably by an altered memory profile. Although clinically well-documented, no specific biomarker exists. This translational study identifies plasminogen activator inhibitor-1 (PAI-1) as a brain risk factor for PTSD, thereby supporting its potential as a blood-derived biomarker. Mice with genetically ablated PAI-1 were protected from developing a PTSD-like memory profile. Conversely, mice exhibiting PTSD-like cognitive impairment showed increased blood PAI-1 levels, correlating with their profile severity. In the brain, PAI-1 levels were specifically increased in the dorsal hippocampus, a key region for cognitive functions and in the etiology of PTSD. Finally, a longitudinal study of soldiers revealed that those developing PTSD symptoms exhibit rising blood PAI-1 levels over a 12-month period. Its significant association with various indicators of PTSD-related psychological distress attests to PAI-1’s potential as a blood biomarker and brain therapeutic target for PTSD.

## Introduction

Post-traumatic stress disorder (PTSD) is a complex stress-related psychiatric condition in response to traumatic life-threatening events. PTSD occurs in 15 to 50% of subjects who have experienced or witnessed of a traumatic stress situation in which their physical and/or psychological integrity was threatened. War situations, violence (including domestic violence), sexual assault, accidents or natural and man-made disasters are common causes of PTSD, leading to severe social, occupational, and interpersonal disturbances. One of the hallmarks of PTSD is a qualitative and paradoxical alteration of memory that involves both hypermnesia of the event based on certain salient elements, although not predictive of the trauma (i.e., sudden resurgence of intrusive images and negative thoughts) and contextual amnesia associated with the traumatic event (i.e., forgetting of the context)^1,2^. Having focused their attention on these salient stimuli, as opposed to the context of the trauma, PTSD subjects subsequently exhibit maladaptive fear responses to these stimuli in non-traumatic contexts. This state leads to maladaptive fear reactions and is associated with the development of behavioral and comorbidities disorders such as depression, early dementia, hyperarousal, sleep disorders, addictions and also suicides^1^.

Although PTSD symptoms are well-documented clinically, the underlying neurobiological mechanisms remain poorly understood. As a result, there are currently no specific biomarker or effective pharmacological treatment available. Treatments using non-selective drugs (e.g., SSRIs, antipsychotics, anticonvulsants, and anti-adrenergic drugs) have often shown limited efficacy and are even less efficient than trauma-focused psychotherapy^3–5^. Given the current geopolitical and societal context, PTSD has emerged as a major public health issue^6,7^. Therefore, identification of blood-derived biomarkers is needed for the pre/post symptomatic diagnosis of this pathology, for which a late diagnosis may lead to chronic PTSD^8^. Currently, PTSD is diagnosed primarily based on clinical criteria, but many factors limit the standardization of the diagnosis. Indeed, clinical evaluations rely heavily on subjective interpretation, which can lead to diagnostic variability between different clinicians. This subjectivity can result in the underdiagnosis or misdiagnosis of PTSD. Second, access to specialized centers for clinical evaluation may be limited in certain countries. This can lead to delays in diagnosis when timely intervention is crucial for improving outcomes. Third, clinical evaluations often focus on symptomatology and may not fully capture the complexity of PTSD, including underlying biological mechanisms and individual variability in response to trauma. Consequently, adopting a more comprehensive evaluation approach grounded in objective measures is essential to substantially improve diagnostic accuracy.

The pathophysiology of PTSD involves intricate neurobiological alterations, including the dysregulation of the hypothalamic-pituitary-adrenal (HPA) axis^9,10^, changes in neurotransmitter systems^11,12^, and structural and functional alterations in key brain regions such as the amygdala, the prefrontal cortex, and notably the hippocampus (Hpc)^13,14^. Evidence was provided that glucocorticoid (GC) hormones (<u>Cort</u>: <u>cort</u>icosterone in rodents, <u>cort</u>isol in humans), one of the main biological responses to stress, and functional alterations of its cognate receptor; glucocorticoid receptor (GR) may play a key role in the onset of PTSD^15–17^. Thus, uncovering the role of specific effectors regulating stress responses and cognition is crucial to understand the neurobiological mechanisms underlying PTSD. Among these, plasminogen activator inhibitor-1 (PAI-1) has emerged as a key mediator in stress-related processes^18,19^. In high stress condition in mice, we have previously demonstrated that GC signaling up-regulate PAI-1 protein in the dorsal Hpc (dHpc)^18,19^, a key structure involved in memory processing^20,21^ and in the etiology of PTSD^22–26^. Stress-induced PAI-1 up-regulation, by disrupting the tissue plasminogen activator (tPA) activity essential for normal memory processing, was shown to trigger PTSD-like fear memory, similar to those observed in PTSD subjects^18,19^. These findings were obtained using an original mouse model, combining contextual fear conditioning and a post-conditioning injection of CORT^27^, which has high translational value as it precisely recapitulates both core components of PTSD memory - emotional hypermnesia and contextual amnesia - distinguishing pathological memory from normal fear memory, thus closely mirroring the dissociation seen in human PTSD memory processing^11^.

The present study explores the emerging role of PAI-1 and its potential involvement in the neurobiological basis of PTSD using a multifaceted mouse-to-human strategy. This strategy combines molecular and cellular tools together with genetically PAI-1 deficient mice and the mouse model that mimics paradoxical memory components of PTSD. In addition, we conducted a longitudinal study on a cohort of soldiers, including biological and psychological outcomes. We showed elevated blood levels of PAI-1 in groups with PTSD-like impairments in mice or a high level of PTSD symptoms in human, as well as a specific increase of PAI-1 protein in the mouse dHpc. Our data highlight the potential of PAI-1 as a promising biomarker of PTSD, causally linked both to PTSD-like memory formation in mice and correlated with markers of psychological distress, such as perceived stress, anxiety, depression and suicidal thoughts that characterize PTSD in humans.

## Results

### Plasma and hippocampal PAI-1 protein and gene encoding (*Serpine1*) expression in response to acute restraint stress

Our previous work has shown that restraint stress, through the release of GC (corticosterone or Cort), enhanced PAI-1 protein expression in the mouse dHpc^18^. This transcriptional modulation relies notably on the presence of GC-responsive elements (GRE) in the promotor region of *Serpine1* encoding PAI-1^29^. Yet the subregion localization of stress effects on hippocampal PAI-1 expression have not been identified. This first series of experiments aimed at characterizing the stress response in PAI-1 deficient mice (PAI-1^-/-^) and also to look at the PAI-1 expression profile in the dHpc of wild-type littermate mice (PAI-1^+/+^) in response to stress. As no significant differences were observed between male and female mice in any of the measured outcomes, data from both sexes were combined for all analyses. Expanding on these prior findings^18^, we show that a 3-hour restraint stress increased Cort levels in plasma samples of both genotypes (**Fig. 1A left panel**, two-way ANOVA; stress effect: F(1, 46)=38.13, *p=1.6.10^-7^;* genotype effect: F(1, 46)=0.00865, *p=0.930,* no interaction), while PAI-1 protein level was detected and increased by stress only in PAI-1^+/+^ mice (**Fig. 1A right panel**, two-way ANOVA; stress effect: F(1, 26)=18.32, *p=0.0002*; genotype effect: F(1, 26)=133,6, *p=1.10^-^* ^11^; interaction: F(1, 26)=18.32, *p=0.0002*). These data show that mice lacking PAI-1 (PAI-1^-/-^) do not exhibit altered peripheral reactivity to stress, as the increase in Cort levels in response to restraint stress was similar to PAI-1^+/+^ mice (**Fig. 1A left panel**). In line with these results, qPCR assays show that expression of *Serpine1* encoding PAI-1 protein increased in the Hpc after stress in PAI-1^+/+^ mice (**Fig. 1B left panel**, t-test; stress effect: *p=0.00003*) and that its expression is absent in the Hpc of mutant mice as expected (**Fig. 1B left panel)**. We also found that restraint stress reduces hippocampal expression levels of glucocorticoid receptor (GR)-encoding *Nr3c1* in mice of both genotypes. These results confirm the efficacy of the Cort inhibitory feedback loop on GR receptor expression, and also that central reactivity to stress is unaffected in PAI-1^-/-^ mice (**Fig. 1B right panel**, two-way ANOVA; stress effect: F(1, 20)=19.71, *p=0,0003;* genotype effect: F(1, 20)=1.422, *p=0.25*, no interaction). Finally, single molecule fluorescent *in situ* hybridization (RNAscope) analysis revealed that although *Serpine1* is expressed in the CA1, CA3 and dentate gyrus (DG) subregions of the dHpc, the highest expression level was found in the stratum radiatum (sr) of CA1, irrespective of the treatment (Control *vs.* Stress). By assessing mRNA levels in dots/mm^2^, we demonstrated a significant increase in *Serpine1* expression in the CA1 and DG of stressed mice as compared to PAI-1^+/+^ control mice (**Fig. 1C**, CA1 t-test *p=0.0463*; CA3 t-test *p=0.2225*; DG t-test *p=0.0140*). Also, as expected, there was a clear absence of *Serpine1* expression in knockout tissues, confirmed by fluorescent RNAscope hippocampal staining (**Supplementary Fig. S1**). Taken together, these data confirm that PAI-1 protein is reactive to stress, with a concomitant increase in PAI-1 in both plasma and hippocampus after restraint stress. Furthermore, our findings also demonstrate that PAI-1^-/-^ mutant mice displayed a total absence of PAI-1 (protein and mRNA) together with a stress reactivity comparable to that of control mice, thereby validating their use for next experiments.

**Fig. 1.**
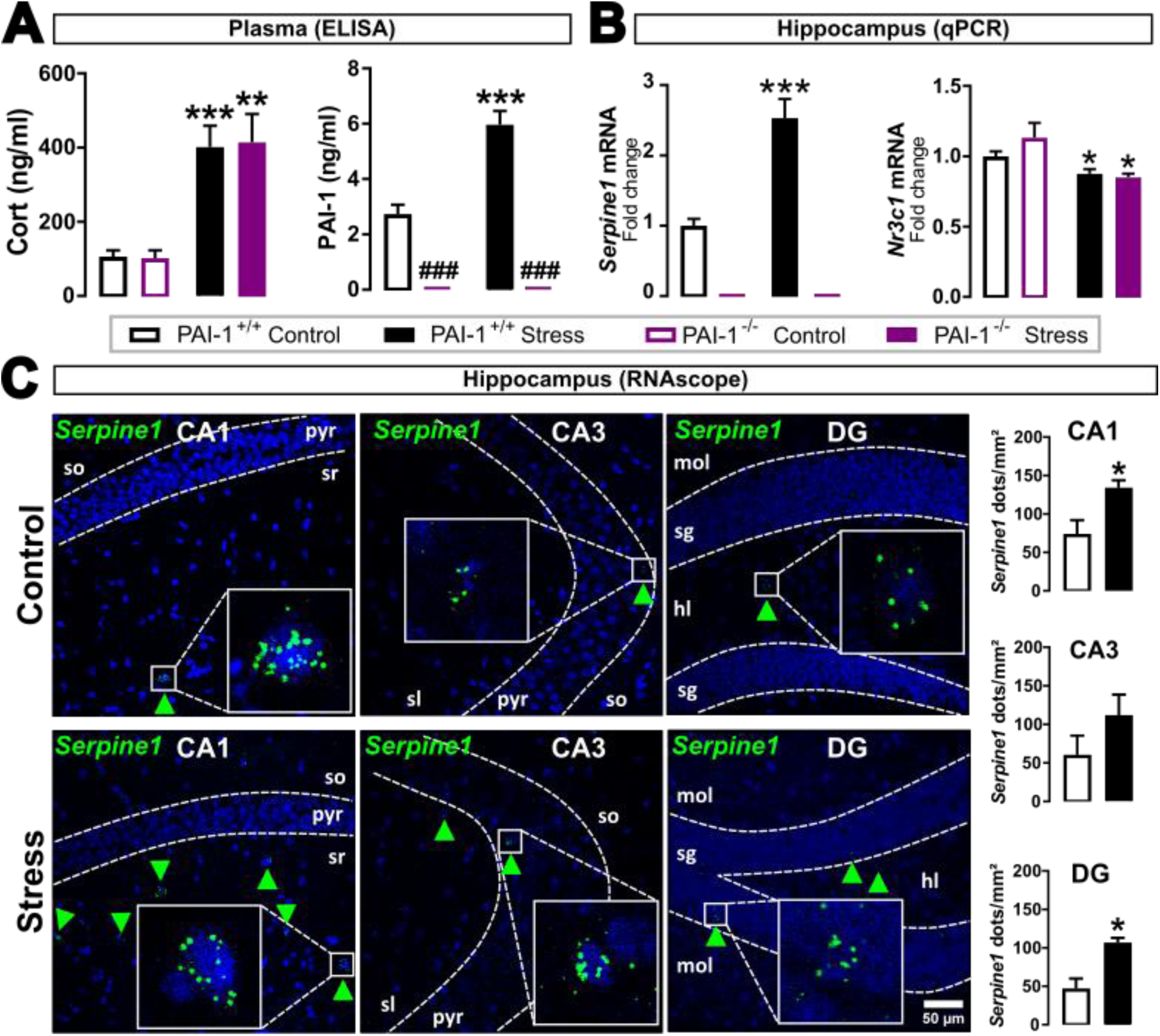
Comparison of the HPA axis responses and PAI-1 levels in plasma and hippocampus in control (PAI-1^+/+^) and PAI-1-deficient (PAI-1^-/-^) male and female mice in response to 3-hour restraint stress. (A) Cort and PAI-1 levels in plasma measured by ELISA in control (WT PAI-1^+/+^) and mutated (KO PAI-1^-/-^) mice under basal conditions (Cort level in WT, n=13; in KO, n=12; PAI-1 level in WT, n=9; in KO, n=6) and in response to restraint stress (Cort level in WT, n=13; in KO, n=12; PAI-1 level in WT, n=9; in KO, n=6). (B) *Serpine1* encoding PAI-1 and *Nr3c1* encoding GR expression levels in the hippocampus (Hpc) measured by qPCR in control and stressed PAI-1^+/+^ (n=7/group) and PAI-1^-/-^ (n=5/group) mice. (C) Confocal images of coronal brain sections of CA1, CA3 and dentate gyrus (DG) of the dorsal Hpc of control (n=4) and stressed (n=3) PAI-1^+/+^ mice showing the distribution of *Serpine1* encoding PAI-1 (green) detected by single-molecular fluorescent *in situ* hybridization (RNAscope). Inserts show high magnification of *Serpine1* positive cells and dots quantification in these sub-regions. Slides were counterstained with DAPI (blue). <u>Pyr:</u> pyramidal cell layer; <u>so</u>: stratum oriens; <u>sr</u>: stratum radiatum; <u>sl</u>: stratum lucidum; <u>mol</u>: molecular layer of the DG; <u>sg</u>: stratum granulosum; <u>hl</u>: hilus of the DG. Data analyzed using two-way ANOVA followed by Fisher LSD post-hoc multiple comparison (A, B) or unpaired t-test (C). Bar graphs represent mean or fold change value of the group in comparison to the PAI-1^+/+^ control group ± sem. \**p<0.05*, \*\**p<0.01* and \*\*\**p<0.001* for stress effect and ^###^*p<0.001* for genotype effect.

### Full deletion of *Serpine1* encoding PAI-1 does not affect behavior under physiological condition in mice

In the next series of experiments, we investigated in detail the behavioral profile of mutant PAI-1^-/-^ compared to control PAI-1^+/+^ mice. First, we measured their basal activity using two behavioral devices. Actimetry for 48 hours was used to check for deficits in circadian rhythm activity, and the open field (OF) test for 30 minutes measured locomotor response to a novel environment. In both tests, we observed no differences between PAI-1^-/-^ and control PAI-1^+/+^ mice. Both genotypes display similar increases in nocturnal activity compared to diurnal activity (**Fig. 2A left panel** <u>actimetry</u>, time effect: F(23, 644)=20.48, *p<0.0001*; genotype effect: F(1, 28)=0.00375, *p=0.9516*; no interaction), as well as similar recovery of activity over time in response to novelty (**Fig. 2A right panel** <u>open-field</u>, time effect: F(5, 195)=43.88, *p<0.0001*; genotype effect: F(1, 39)=0.05687, *p=0.8128*; no interaction). We then investigated whether different types of memory were affected following PAI-1 deletion. We assessed working memory by measuring alternation in the Y-maze test, spatial memory using a spatial Y-maze task and object recognition using the novel object recognition (NOR) test. In the Y-maze test, we found that both PAI-1^+/+^ and PAI-1^-/-^ mice exhibited correct memory performance in alternation, as evidenced by significantly above-chance exploration of the three arms (**Fig. 2B left panel**, one sample t-test PAI-1^+/+^ *p=8.10^-7^* and PAI-1^-/-^ *p=4.10^-5^*) and correct recognition memory, showing the ability to distinguish between a previously visited arm (familiar arm or FA) and a new arm (NA) using spatial cues (**Fig. 2B right panel**, one sample t-test PAI-1^+/+^ *p=3.10^-^*^10^ and PAI-1^-/-^ *p=1.10^-7^*). Similarly, in NOR test, both PAI-1^+/+^ and PAI-1^-/-^ mice were able to distinguish between familiar (FO) and novel object (NO) (**Fig. 2C**, one sample t-test PAI-1^+/+^ *p=8.10^-7^* and PAI-1^-/-^ *p=0.0004*) with no effect of PAI-1 genetic deletion. Also, fear-related associative memories were measured using two well-known protocols of Fear Conditioning (FC) procedures: cued and contextual FC. We showed that there was no effect of the genotype on memory performance in terms of freezing behavior during habituation, conditioning, re-exposure and extinction phases and that both genotypes were able to extinguish previously acquired fear (**Fig. 2D**, <u>cued FC</u>: time effect: F(5,135)=47.98, *p<0.0001*; genotype effect: F(1,27)=0.01385, *p=0.9072*; no interaction and <u>contextual FC</u>: time effect: F(4,108)=62.71, *p<0.0001*; genotype effect: F (1,27)=0.1560, *p=0,6960*; no interaction). Finally, we assessed reactivity to stress using stress-induced hyperthermia (SIH) (**Fig. 2E**), depression-like behaviors with the forced swim test (FST) and the sucrose preference (SP) test (**Fig. 2F**), and anxiety-like behaviors with the elevated plus maze (EPM), open field (OF) and light-dark box (LD) tests (**Fig. 2G**). Altogether, no differences between genotypes/gender were observed in any of these behavioral paradigms, indicating that complete deletion of PAI-1 does not affect behavior under physiological conditions in mice.

**Fig. 2.**
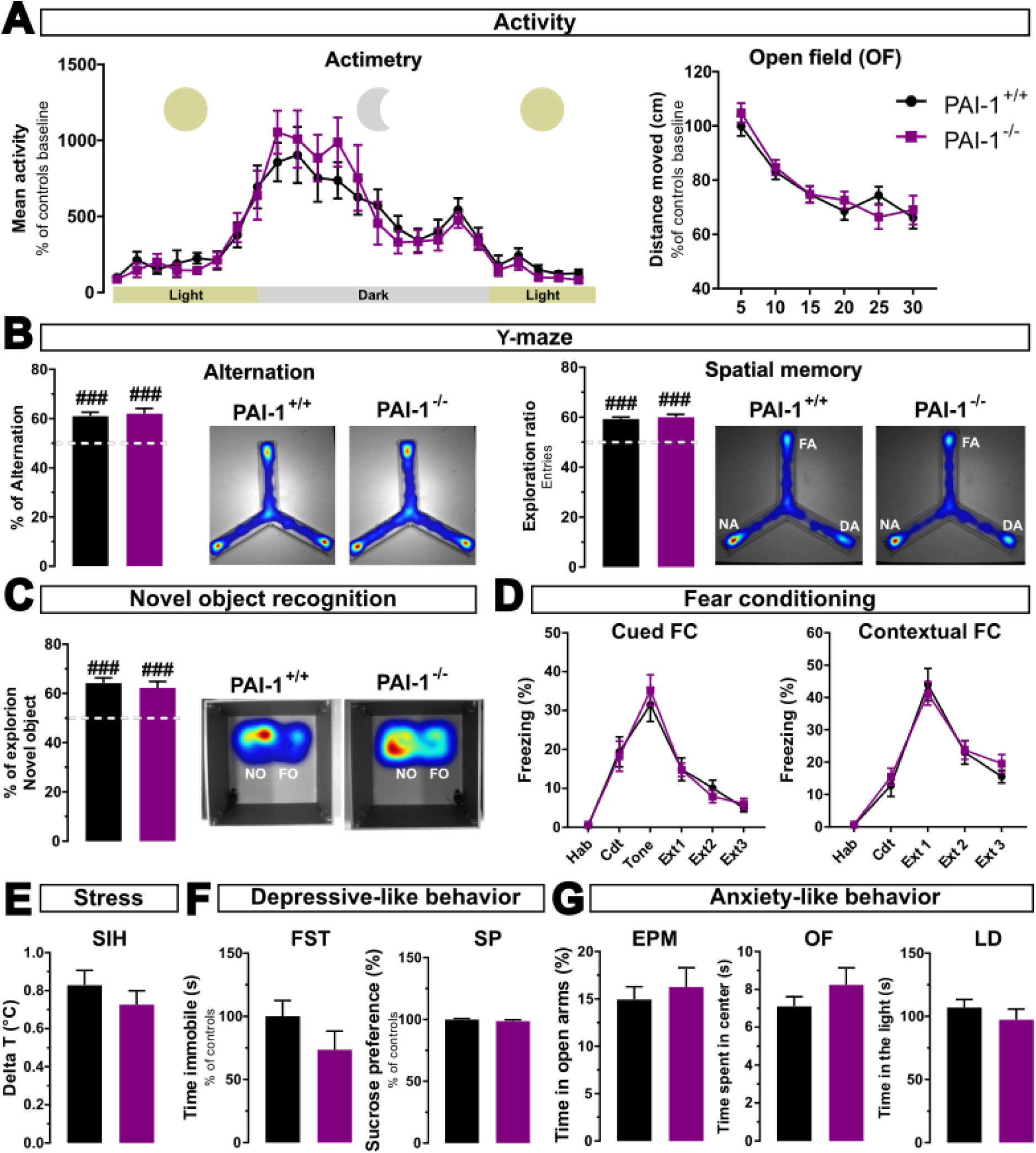
PAI-1-deficient (PAI-1^-/-^) male and female mice show no differences in several behavioral tasks under physiological conditions compared to control littermate (PAI-1^+/+^) mice. **(A)** Mean activity of PAI-1^+/+^ (actimetry n=14, OF n=18; black squares) and PAI-1^-/-^ (actimetry n=16, OF n=23, purple squares) mice for 2 days in actimetry cages using beam detection (left panel) and distance traveled during 30-minute open field (OF) activity measurement (right panel). **(B)** Y-maze alternation (% of alternance, left panel) and spatial memory tests (exploration ratio of entries, right panel) of PAI-1^+/+^ (n=23) and PAI-1^-/-^ (n=15) mice. White dash bars represent chance level and representative heatmap of both genotypes are presented (FA: familiar arm, DA: departure arm, NA: novel arm). **(C)** Novel object recognition memory test of PAI-1^+/+^ (n=23) and PAI-1^-/-^ (n=15) mice. White dash bar represents chance level and representative heatmap of both genotypes are presented (FO: familiar object, NO: novel object). **(D)** Fear conditioning (FC) with cued (left panel) or contextual protocols (right panel) of PAI-1^+/+^ (Cue FC n=13, Cx FC n=14) and PAI-1^-/-^ (Cue FC n=16, Cx FC n=15) mice. Freezing levels during the different days of protocol are presented (Hab: habituation, Cdt: conditioning, Tone: tone test for cued FC, Ext: extinction). **(E)** Stress-induced hyperthermia as an index of stress reactivity of PAI-1^+/+^ (n=27) and PAI-1^-/-^ (n=16) mice. **(G)** Depressive-like behavior of PAI-1^+/+^ (FST n=20, SP n=24) and PAI-1^-/-^ (FST n=14, SP=15) mice. **(F)** Anxiety-like behavior of PAI-1^+/+^ (EPM n=23, OF n=20 and LD n=26) and PAI-1^-/-^ (EPM n=15, OF n=20 and LD n=15) mice. SIH: stress induced hyperthermia, FST: forced swim test, SP: sucrose preference, EPM: elevated plus maze, OF: open field, LD: light/dark box. **(A, D)** Data analyzed using two-way ANOVA repeated measures followed by Fisher LSD post-hoc multiple comparison. **(B, C, E, F, G)** unpaired t-test. For each graph mean ± sem is shown. ### *p<0.001* different from chance, for comparison to chance level unpaired t-test was used.

### Full deletion of *Serpine1* encoding PAI-1 protects mice from PTSD-like memory impairment

Our previous report identified PAI-1 as a critical molecular switch between moderate stress and adaptive memory processes, and PTSD-like maladaptive fear memory, since PAI-1 directly injected into the mouse dHpc was sufficient to induce a PTSD-like memory profile in a condition of fear conditioning known to normally induce an adaptive fear memory^18^. Here, we investigated whether PAI-1 deletion could prevent the onset of the PTSD-like memory profile. To address this issue, we used a well-established PTSD-like mouse model, which combines contextual FC and a post-conditioning injection of Cort^27^. This model successfully recapitulates two memory features of PTSD: emotional hypermnesia for a salient cue (tone), which is non-predictive of the unconditioned stimulus (US) and contextual amnesia for contextual cues from the conditioning chamber (context = Cx)^1,28^. As such, these animals, like PTSD patients, lose their ability to restrict fear to the right situation or cue. Contextual FC is immediately followed by an intraperitoneal injection of Cort (2.5 mg/kg) or NaCl (control vehicle) and re-exposure to tone and Cx was performed on days 3 and 4, respectively (**Fig. 3A**). As expected, NaCl-injected control PAI-1^+/+^ mice showed no fear response (no freezing) to the irrelevant (non-predictive) tone (**Fig. 3B left panel**). However, when PAI-1^+/+^ mice were injected with Cort immediately after conditioning, the PTSD-like memory profile appeared, since they displayed an abnormally high fear response to the salient tone, although non-predictive, characteristic of the PTSD-like memory profile (**Fig. 3B left panel**). In contrast, this maladaptive PTSD-like memory profile does not appear in PAI-1 mutant mice injected with Cort which did not differ from their NaCl-injected control mutant mice (**Fig. 3B right panel**). These results are illustrated by the tone ratio in the 4 groups of mice (**Fig. 3C**, two-way ANOVA; <u>treatment effect</u>: F(1, 110)=2.682, *p=0.1043*; <u>genotype effect</u>: F(1, 110)=3.692, *p=0.0573*; <u>interaction</u>: F(1, 110)=6.493, *p=0.0122*). During contextual re-exposure, we did not observe significant differences over the 6-minute period (**Fig. 3D**, two-way ANOVA for the 6-minutes period; time effect: F(1.514,166.5)=160, *p<0.0001*; genotype effect: F(3,110)=0.9461, *p=0.4210*; interaction: F(6,220)=2.138, *p=0.0502*). However, by looking specifically at freezing levels during the first two minutes of the context test (Cx 0-2), we identified contextual amnesia as seen by a decreased freezing response to the context in PAI-1^+/+^ mice injected with Cort compared to NaCl-injected mice (t-test, PAI-1^+/+^ NaCl vs Cort, *p=0.012*). Here again, no maladaptive fear response to context following Cort injection was observed in PAI-1^-/-^ mutant mice (**Fig. 3D right panel**, two-way ANOVA for Cx 0-2; <u>treatment effect</u>: F(1, 110)=1.718, *p=0.193*; <u>genotype effect</u>: F(1, 110)=0.673, *p=0.414*; <u>interaction</u>: F(1, 110)=2.898, *p=0.092*).

**Fig. 3.**
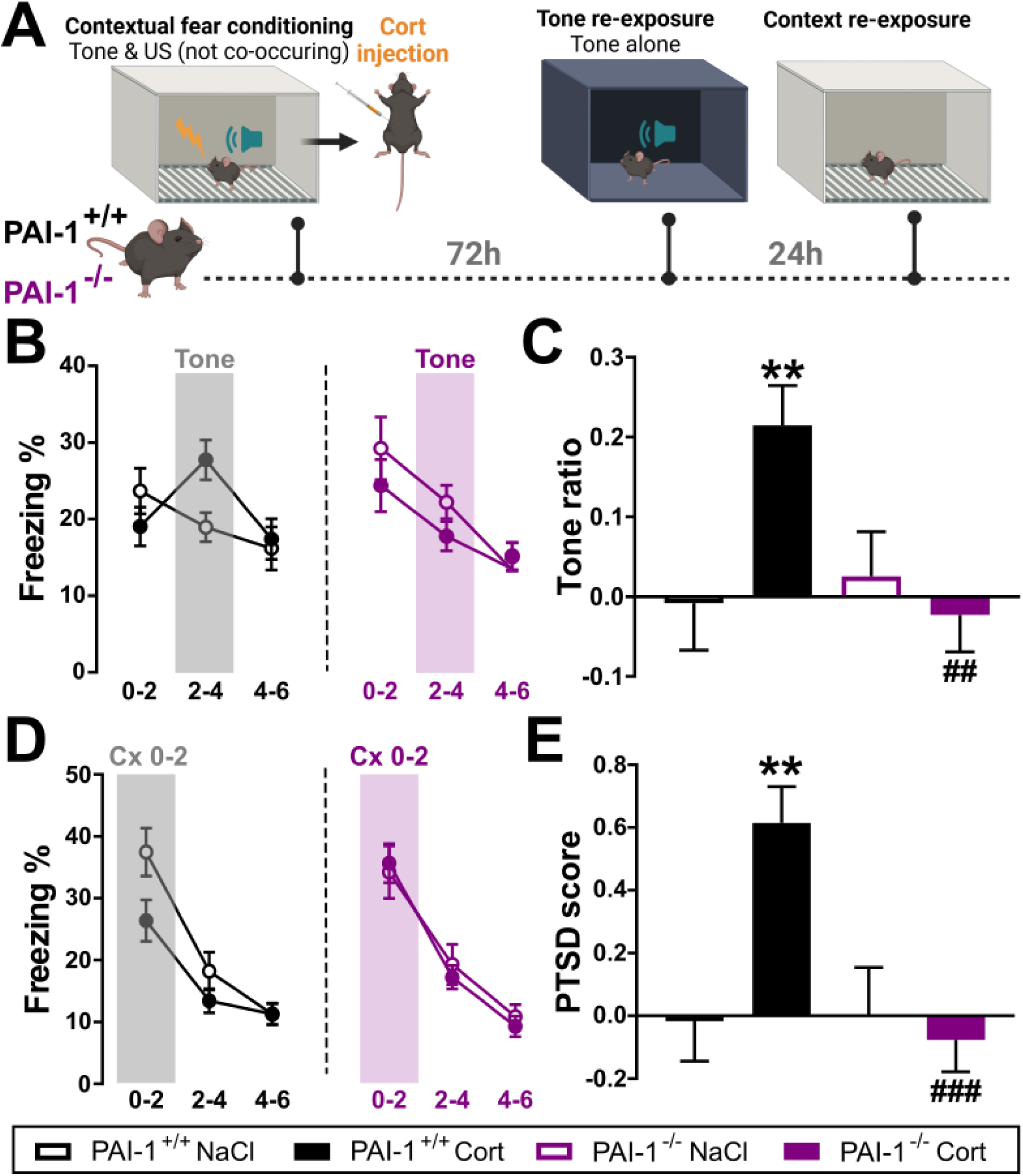
PAI-1-deficient male and female mice (PAI-1^-/-^) do not develop a PTSD-like memory profile. **(A)** Schematic of the protocol timeline represented by the fear conditioning (FC) protocol with unpaired presentations of two tones and two shocks (unconditioned stimulus or US) followed by an injection of corticosterone (Cort, 2.5 mg/kg) or NaCl (control vehicle) then re-exposed to both tone (day 5) and context (day 6). **(B)** Freezing levels during re-exposure to tone after contextual FC followed by injection of either NaCl (empty dots) or Cort (filled dots), represented by 2 min time blocks (0-2, 2-4 and 4-6 min) in control littermate (PAI-1^+/+^) and deficient (PAI-1^-/-^) mice. Tone is presented during the 2-4 min time block. **(C)** Response to the tone represented as a tone ratio, calculated with the following formula using freezing level during time blocks: tone ratio = (2-4 - ((0-2 + 4-6)/2)) / (2-4 + ((0-2 + 4-6)/2)). **(D)** Freezing levels during re-exposure to context (Cx) after contextual FC followed by injection of either NaCl (empty dots) or Cort (filled dots), represented by 2 min time blocks (0-2, 2-4 and 4-6 min) in control littermate (PAI-1^+/+^) and deficient (PAI-1^-/-^) mice. Typically, differences between mice treated with NaCl and Cort are seen in the first block (Cx 0-2). **(E)** Calculated PTSD score based on the average of two z-scores combining the fear responses to the tone and the context. Groups: PAI-1^+/+^ NaCl n=29 & Cort n=27 and PAI-1^-/-^ NaCl n=28 & Cort n=30. Data analyzed using two-way ANOVA with repeated **(B, D)** or not repeated **(C, E)** measures followed by Fisher LSD post-hoc multiple comparison. For each graph mean ± sem is shown. \*\**p<0.01* for treatment effect and ## *p<0.01* and ### *p<0.001* for genotype effect.

Finally, to provide a global profile of PTSD-like memory that encompasses both tone and context responses, we computed a PTSD score based on tone ratio and freezing during Cx 0-2 respectively for re-exposure to tone and context. Using this score, we confirmed that while PAI-1^+/+^ mice injected with Cort clearly present a PTSD-like memory profile compared to NaCl-injected mice, whereas PAI-1^-/-^ mutant mice do not (**Fig. 3E**, two-way ANOVA for PTSD score; <u>treatment effect</u>: F(1, 110)=4.262, *p=0.041*; <u>genotype effect</u>: F(1, 110)=6.995, *p=0.009*; <u>interaction</u>: F(1, 110)=7.023, *p=0.0092*). Altogether, these results show that PAI-1 full deletion protects from developing a paradoxical PTSD-like memory profile in mice of both genders.

### Increased PAI-1 blood level in mice with PTSD-like fear memory impairment

In these series of experiments, we investigated the potential of blood PAI-1 protein as a biomarker and brain risk factor for PTSD-like memory profile in mice. Our hypothesis is that, due to the pleiotropic action of GC on *Serpine1* encoding PAI-1, mice with PTSD-like memory profile should show increased blood levels of PAI-1 in comparison to mice with adaptive fear memory. To address this issue, at the end of the behavioral procedure, we collected blood and brain samples from C57BL/6N mice of both genders having developed a PTSD-like memory profile (i.e., injected with Cort) and those that displayed an adaptive conditioned fear memory (i.e., injected with NaCl) (**Fig. 4A**). As expected, Cort-injected mice exhibited a PTSD-like memory profile, with hypermnesia, as shown by their strong fear response to the tone (**Fig. 4B-C**, t-test for tone ratio *p=0.0002*), as well as contextual amnesia, as shown by their decreased fear response to the context (**Fig. 4D**, two-way ANOVA for context test; time of the test effect: F(2, 36)=8.228, *p=0.001*; treatment effect: F(1, 18)=7.031, *p=0.016*; interaction: F(18, 36)=3.904, *p=0.0002*). This was also illustrated by their higher PTSD score compared to NaCl-injected mice (**Fig. 4E**, t-test for PTSD score *p=0.0001*). We then measured their plasma PAI-1 levels and found higher circulating PAI-1 levels in Cort-injected mice when compared to NaCl-injected mice (**Fig. 4F left panel**, t-test PAI-1 plasma level *p=0.01*). Interestingly, plasma PAI-1 levels were further correlated with their PTSD scores (**Fig. 4F right panel**, r^2^ = 0.849, *p=0.0011*). Finally, we also performed qPCR analysis of *Serpine1* encoding PAI-1 by collecting the hippocampus dissociated into its dorsal (dHpc) and ventral (vHpc) parts, as well as the medial prefrontal cortex (mPFC) and amygdala (Amg), since these structures are also involved in the etiology of PTSD^13^. Interestingly, we found that the increase in *Serpine1* expression was specifically restricted to the dHpc (**Fig. 4G**, t-test dHpc *p=0.0017*, vHpc *p=0.9032,* mPFC *p=0.8167,* Amg *p=0.7448*). Such increase was also observed in dHpc subregions using RNAscope *in situ* hybridization after restraint stress (**Fig. 1C**). This finding further strengthens the key role of dHpc in PTSD, and in particular in the switch between adaptive and maladaptive (PTSD-like) fear memory^30^, in addition to its well-known role in fear memory processing, contextual learning and stress response regulation^18,30–33^. Altogether, these data in mice support a role for PAI-1 as a blood biomarker and a brain risk factor associated with PTSD-like memory deficits.

**Fig. 4.**
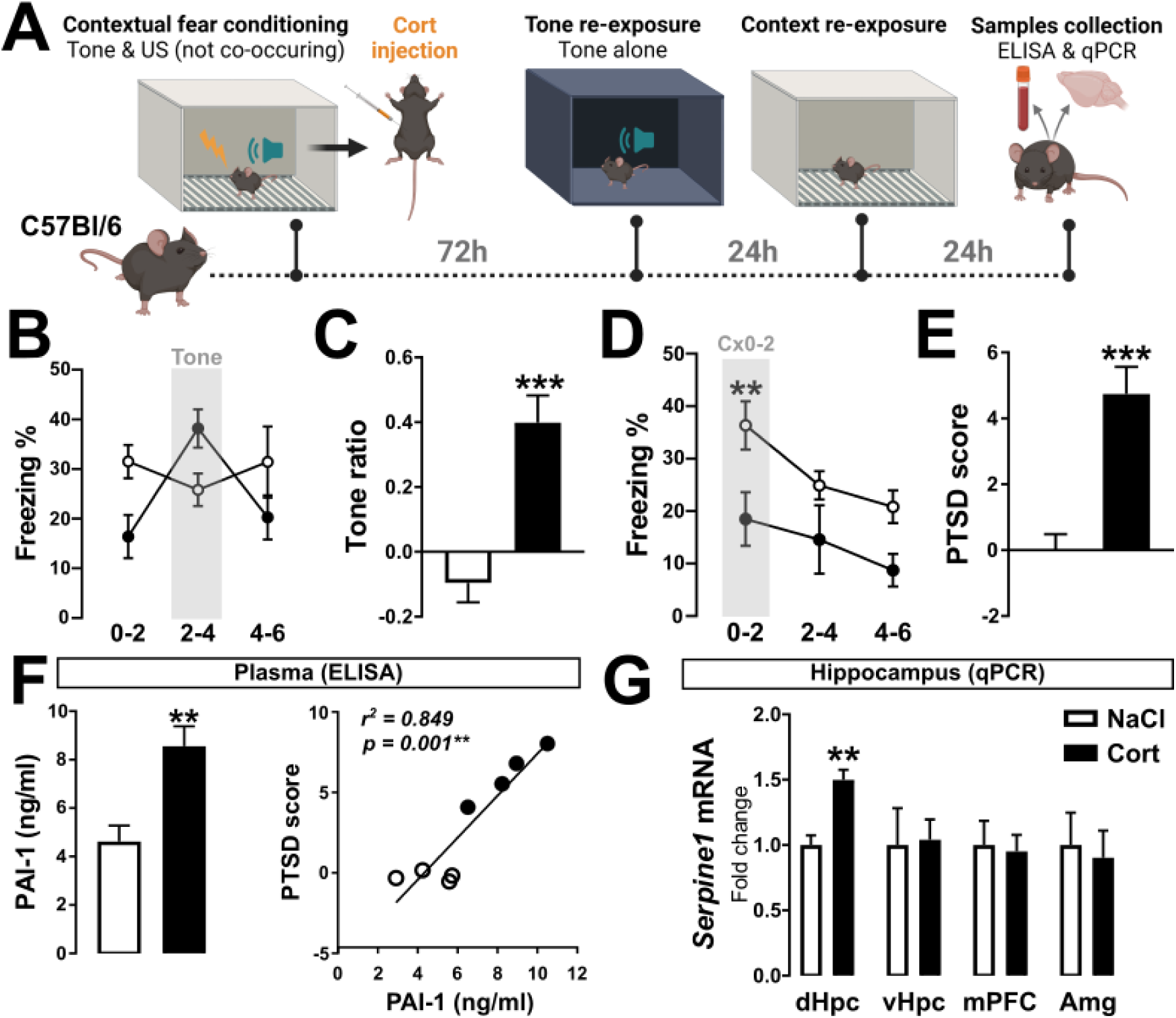
Increase in PAI-1 blood and dorsal hippocampus levels in male and female mice with PTSD-like memory impairment. **(A)** Schematic of the protocol timeline. **(B)** Freezing levels during re-exposure to tone after contextual fear conditioning followed by injection of either NaCl (white) or Cort (2.5 mg/kg; black), represented by 2 min time blocks (0-2, 2-4 and 4-6 min). Tone is presented during the 2-4 min time block. **(C)** Response to the tone represented as a tone ratio. **(D)** Freezing levels during re-exposure to context (Cx) after contextual fear conditioning followed by injection of either NaCl (white) or Cort (2.5 mg/kg, black), represented by 2 min time blocks (0-2, 2-4 and 4-6 min). **(E)** PTSD score calculated using the tone ratio and the freezing during the first two minutes of re-exposure to the context (Cx 0-2). Groups: NaCl n=10 and Cort n=10. **(F)** Plasma PAI-1 levels after induction of PSTD-like memory profile in mice, measured by ELISA (left) and correlation with PTSD score (right) (NaCl n=4 & Cort n=5). **(G)** Expression levels of *Serpine1* encoding PAI-1 in the dorsal and ventral hippocampus (dHpc and vHpc), medial prefrontal cortex (mPFC) and amygdala (Amg) measured by qPCR in mice injected with NaCl (n=5) and Cort (n=5). **(B, D)** Data analyzed using two-way ANOVA for repeated measures followed by Fisher LSD post-hoc multiple comparison, **(C, E, F left, G)** Unpaired t-test and **(F right)** linear regression. For each graph mean ± sem is shown and for linear regression individual data points are presented; \*\**p<0.01* and \*\*\**p<0.001* for treatment effect.

### Increased PAI-1 blood level in human subjects with PTSD symptoms

The previous preclinical findings pointing to similar PAI-1 increase between the periphery and the brain suggest that PAI-1 could represent a valuable biomarker for detecting stress sensitivity, as for quantifying clinical severity and assessing the clinical prognosis of PTSD. To address this pertinent issue, we measured PAI-1 blood levels in a longitudinal cohort of soldiers (n=80) who had been deployed to a war zone for 6 months and who had been subjected to a psychological assessment of their traumatic state by means of questionnaires^34,35^. According to the posttraumatic checklist scale questionnaire (PCL-S), which recapitulates the three main symptoms of PTSD (revival, avoidance and hyperactivity)^36^ and which remains the reference for PTSD symptoms in the French Army, around 20% of soldiers had a high level of PTSD symptoms (PTSD-S) and were classified as being at high risk for PTSD in the near future (**Fig. 5A**). Firstly, we found that pre- and post-mission PAI-1 levels were highly correlated, indicating good biological validity of samples and measurements (**Fig. 5B**). As a result, we compared the difference in PAI-1 levels after and before the mission (ΔPAI-1) to normalize levels for each soldier and focus on the evolution of PAI-1 over time with regard to mission-associated trauma. Data analysis of this longitudinal cohort showed that ΔPAI-1 increased in soldiers with high levels of PTSD symptoms (PTSD-S; PCL-S score > 34), whereas conversely it decreased in control soldiers with low levels of PTSD symptoms (CON, PCL-S score < 34) (**Fig. 5C**, t-test for ΔPAI-1, CON vs PTSD-S, *p=0.002*). Similarly, analysis of the individual population showed that the proportion of soldiers with increasing ΔPAI-1 was higher in the PTSD-S group than in the CON group (increasing red ΔPAI-1 61.5% *vs.* 31.3%, respectively) and vice versa for decreasing (decreasing green ΔPAI-1 38.5% *vs.* 68.7%, respectively) (**Fig. 5D**). Interestingly, by comparing ΔPAI-1 with other psychopathology questionnaire scores, which further consolidate the diagnosis of PTSD, we found that ΔPAI-1 negatively correlates with vertical group cohesion, considered as a protective factor, and positively correlates with known risk factors such perceived stress (Cohen), anxiety and depression (Hospital anxiety and depression scale or HADS) and also suicidal thoughts (extracted from the general health questionnaire or GHQ-28) (**Table 1**). Finally, we investigated the potential of several cut off values of ΔPAI-1 as a diagnostic tool for PTSD using formulas for sensitivity, specificity, predictive positive and negative values (**Table 2**). We found that a cut off value of 30 ng/ml seems to show good specificity (82.1%) and in particular with a high predictive value to screen soldiers with low PSTD symptoms (Negative Predictive Value or NPV=91.7% chance, **Table 2**). In addition, we calculated for each cut off value of ΔPAI-1 the Youden index defined as a unique and quantifiable measure of the overall performance of a diagnostic test^37^. A cut off value of 30 ng/ml gives the highest Youden index (0.44) which is in moderate range performance but commonly observed in psychiatry due to biological complexity of the pathology and the subjectivity of the diagnostic^38^. Taken together, the analysis of a longitudinal cohort of soldiers predisposed to PTSD revealed that those who develop PTSD symptoms exhibited increasing level of blood PAI-1 levels. These results also strongly suggest that elevated PAI-1 may be a risk factor for PSTD and that its evolution over time in people experiencing trauma could predict whether they will develop a PTSD following trauma exposure.

**Fig. 5.**
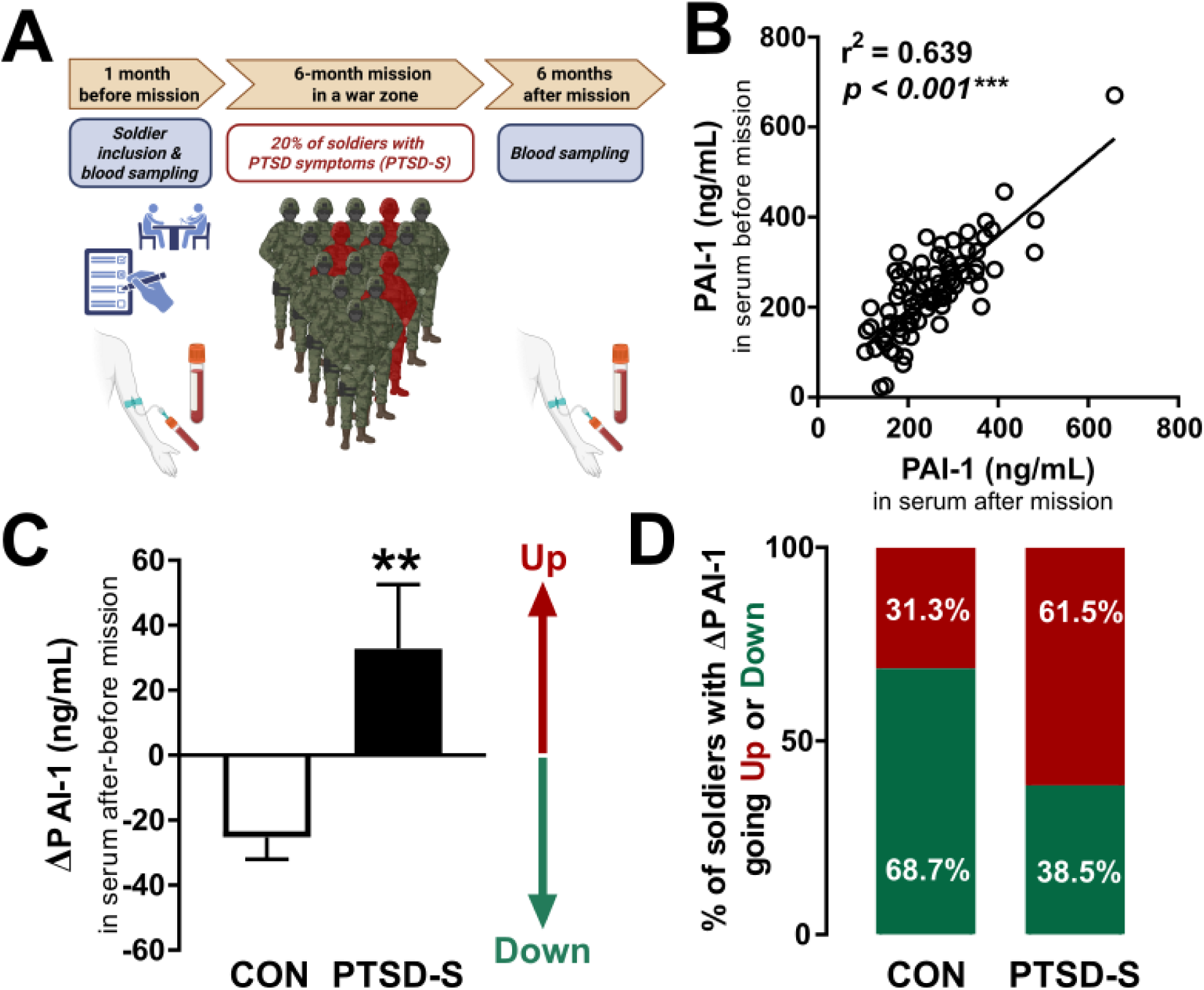
Increase in PAI-1 blood levels over time in male soldiers with PTSD symptoms (PTSD-S). **(A)** Graphical representation of the French soldier’s cohort deployed in Afghanistan (n=80). Blood samples were collected 1 month before and 6 months after their return from a 6-month mission in war zones. The psychological assessment of their traumatic state was carried out using various questionnaires. **(B)** Linear regression of PAI-1 levels, including individual data points, in serum before and after ELISA assays, \*\*\**p<0.001* for linear regression effect. **(C)** PAI-1 levels after – before (ΔPAI-1) in subjects exhibiting low levels of PSTD symptoms (CON, n=67) and subjects with high levels of PTSD symptoms (PTSD-S, n=13). Plotted values are mean ± sem. Unpaired t-test, \*\**p<0.01* for group effects. **(D)** Proportion of soldiers in percentage (%) with increasing (red) or decreasing (green) ΔPAI-1 levels in each group.

**Table 1.**
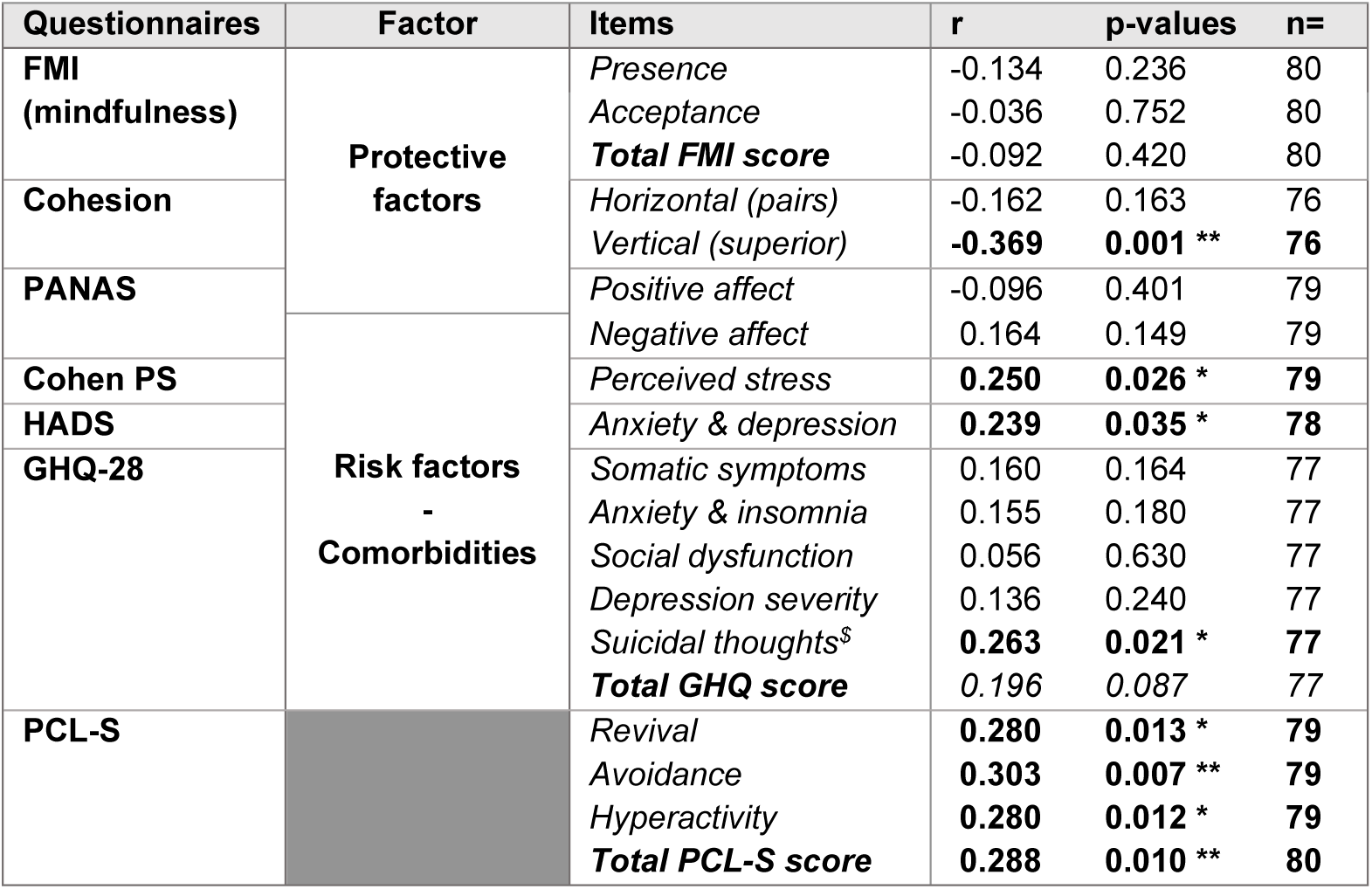
Correlations between ΔPAI-1 and items from various clinical questionnaires. FMI: Freiburg Mindfulness Inventory, PANAS: Positive Affect and Negative Affect Scale, Cohen PS: Cohen Perceived Stress, HADS: Hospital Anxiety and Depression Scale, GHQ-28: General Health Questionnaire, PCL-S: PTSD Checklist Scale. ^$^: correlation with suicidal thoughts was calculated based on questions 24, 25, 27 and 28 of GHQ-28 questionnaire. \**p<0.05* and \*\**p<0.01* for linear regression effect.

**Table 2.** Evaluation of ΔPAI-1 as a diagnostic tool for PTSD symptoms. This table depicts sensitivity and specificity calculation for a potential biomarker. Sensitivity shows the ability of a biomarker to diagnose people with the condition (i.e., percentage of true positives) and specificity to identify those who do not have the condition (i.e., percentage of true negatives). Positive predictive value (PPV) demonstrates the number of true positives in the pool who tested positive and negative predictive value (NPV) the number of true negative in the pool who tested negative. Youden index is used to evaluate the global performance of a biomarker tool for diagnosis based on both sensitivity and specificity.

| Cut off value of ΔPAI-1 (ng/ml) | Sensitivity (%) | Specificity (%) | Positive Predictive Value (PPV in %) | Negative Predictive Value (NPV in %) | Youden Index |
| --- | --- | --- | --- | --- | --- |
| 0 | 61.5 | 68.7 | 27.6 | 90.2 | 0.30 |
| 10 | 61.5 | 71.6 | 29.6 | 90.6 | 0.33 |
| 20 | 61.5 | 80.6 | 38.1 | 91.5 | 0.42 |
| 30 | 61.5 | 82.1 | 40.0 | 91.7 | 0.44 |
| 40 | 53.9 | 85.1 | 41.2 | 90.5 | 0.39 |
| 50 | 38.5 | 92.5 | 50.0 | 88.6 | 0.31 |

## Discussion

A major challenge in psychiatry is to provide an accurate and reliable diagnosis to screen for mental illnesses, so that therapeutic solutions can be considered in the long term. PTSD exemplifies this broader psychiatric challenge of diagnosing a biologically heterogeneous illness using subjectively reported symptoms. The present study shows consistently across both mouse and human species an increase in blood levels of PAI-1 protein under traumatic stress conditions typically encountered by PTSD subjects. This peripheral blood increase in mice and in human was further matched by an increase in PAI-1 specifically in the dHpc of mice. In human these elevated blood PAI-1 levels also showed a strong association with various indicators of psychological distress characteristic of PTSD, including anxiety, depression, perceived stress and suicidal thoughts. These data suggest that PAI-1 has potential as a diagnostic biomarker for this pathology. In addition, PAI-1 protein invalidation in mice was shown to prevent specifically the onset of PTSD-like memory deficits without affecting learning under physiological conditions, suggesting that pharmacological inhibition of PAI-1 could also provide a therapeutic avenue for this pathology.

### Blood-derived PAI-1 as a potential biomarker for PTSD

To the best of our knowledge, there are no circulating biomarkers for the diagnosis of PTSD. This is a real issue, as the absence of objective measures means that the diagnosis of PTSD relies predominantly on self-reported symptoms and interviews with clinicians. In addition, overlapping with other disorders, such as depression and anxiety, as well as cultural and contextual variability influence on how trauma is expressed, are likely to complicate the diagnosis, thus leading to misdiagnosis, while failing to stem the fear and stigma conveyed by this disease^39,40^. PTSD complications are frequent, since first symptoms often emerge months post-trauma, suggesting that timely access to intervention is critical. High-risk professions (e.g., firefighters, police and military forces) are particularly targeted, for whom self-stigmatization of the disease and low medical care delay the diagnosis, leading to chronic PTSD^8^. For civilians, the current societal context (wars, terrorist attacks, post-Covid-19 complications^41^) suggests a future increase in PTSD prevalence^42,43^. Our findings have associated preclinical and clinical validation studies, using both mouse model with PTSD-like memory impairment and human subjects with PTSD symptoms, to allow identification of blood PAI-1 as potential valuable biomarker for PTSD. In our preclinical study on mice, we first showed that in response to stress and after the induction of a PTSD-like memory profile, PAI-1 was significantly increased in the blood. We also found a significant correlation between stress-induced PAI-1 concentrations in the blood and the severity of PTSD-like memory impairments (i.e., PTSD score) in mouse. Lastly, our clinical studies demonstrated the efficacy and reliability of our PAI-1 biomarker-based diagnostic approach, showcasing a relatively high sensitivity (above 60%) and high specificity (above 80%) in identifying PTSD cases. Consequently, PAI-1 measurement assay could be extremely useful as a screening tool, particularly for ruling out the diagnosis of PTSD, due to its high negative predictive value (above 90%)^44^. The negative predictive value indicates the probability that a person with a negative result is actually disease-free. The Youden index, used to evaluate effectiveness of a diagnostic test by measuring the ability to distinguish between true positives and true negatives, gave a score of 0.44. Although, this score is in moderate performance range (from 0: the test performs no better than chance to 1 the test has perfect sensitivity and specificity), it is typical in psychiatry due to the biological complexity of the pathology^38^. To improve diagnostic accuracy and complement clinical assessments, a possible option might be to combine PAI-1 assays with neuroimaging studies, which have shown reduced hippocampal volume and amygdala hyperactivity in PTSD patients using fMRI^23,45–47^. Alternatively, another option would be to pair PAI-1 measurements with analyses of genetic polymorphisms encoding human *SERPINE1*. Indeed, several variations of polymorphisms in PAI-1 promoter and 3′-UTR regions were shown to increase PAI-1 expression levels and have been associated with risk of cardiovascular pathologies, notably thrombotic events^48,49^. The variability within the human cohort could be due to these polymorphisms in human *SERPINE1*^48,50^. Although, there are no direct evidence linking these genetic polymorphisms to PTSD, this potential association deserves further investigation.

### PAI-1 as a brain risk factor for PTSD

In this study, we also measured PAI-1 in the Hpc in stress condition and after the induction of a PTSD-like memory profile in mice. Notably, we showed that PAI-1 gene expression level was up-regulated specifically in the dorsal part of the Hpc in mice with PTSD-like memory profile. Together with the results from our previous study^18^, these data further strengthens the key role of this brain region in contextual memory in mice^51,52^, known to be impaired in our PTSD model^27,30^. Interestingly, this brain region is playing a central role in contextual memory processing that are also affected in PTSD patients^31,32^ and brain imaging studies consistently highlighted a link between PTSD and hippocampal atrophy^23,24,26^, although, the causal link remains debated. In agreement with these results, independently of PAI-1 increase, the dHpc was shown to be a sensitive brain structure, particularly vulnerable to cerebral injury such as head trauma, ischemia, stroke, status epilepticus and Alzheimer disease^53,54^ and is known to be more prone to dysfunction under pathological conditions^55^ likely due to its molecular and structural susceptibility profile^56^. Alternatively, it may also be that the selective expression of PAI-1 and the underlying PAI-1-mediated pro-apoptotic signaling cascade within this brain structure account for the vulnerability of dHpc in this pathology. Indeed, PAI-1 upregulation may influence synaptic plasticity^57^ in this region through the inhibition of the tPA/BDNF pathway^58–60^, inducing neural cell death^61^ thereby affecting the cognitive outcomes associated with trauma exposure observed in individuals with PTSD. The hypothesis of a dHpc-dependent transcriptional increase in *Serpine1* encoding PAI-1 is also supported by evidence that the dHpc exhibits the highest concentration of GR among brain regions^62,63^ and that *Serpine1* is upregulated by stress^64,65^ as it contains GRE in its promotor region^29^. Lastly, evidence has also shown that PAI-1 may originate from endothelial cells^66^, which would explain both its peripheral expression and that of dHpc. Indeed, its expression pattern mainly in the CA1 stratum radiatum (sr) is consistent with this hypothesis as this subregion is rich in blood vessels and consequently in endothelial cells. In response to increased GC concentrations following traumatic confrontation, endothelial cells could then induce an increase in PAI-1 and its local extravasation via an enhancement of the permeability of the blood-brain barrier^67^. This is particularly relevant considering that dHpc is particularly sensitive to blood flow variations^68^ in alignment with its increased vulnerability to traumatic stress exposure.

### PAI-1 as a potential therapeutic target to treat PTSD

Our results using genetically invalidated mice for *Serpine1* encoding PAI-1 showed that PAI-1 has a major role in the PTSD-like memory process, as these mice were protected from the onset of the paradoxical PTSD-like memory profile. PAI-1 is a serine protease inhibitor that plays a pivotal role in the regulation of the fibrinolytic system^69^ and is implicated in several pathological conditions, including cardiovascular diseases^70,71^, metabolic disorders^72,73^, fibrosis^74,75^, cancer^69,76^ and infectious diseases^77,78^, as well as in various neurobiological processes, including neuroinflammation^79–82^ and in stress-related conditions^18,19^. Therefore, its invalidation could be expected to have detrimental effects. Evidence in humans showed that the absence of PAI-1 has a paradoxical impact on cardiovascular and hemorrhagic risks, either beneficial or deleterious depending on the pathological context and gender. However, this is restricted to abnormal bleeding and is only observed after trauma or surgery in homozygous affected individuals^83–86^. Our data on PAI-1 deficient mice showed that they were viable and fertile, with no hemorrhagic or cardiovascular abnormalities and their circadian rhythmic activity was unaffected. In addition, their response to acute restraint stress was within a normal range, with a properly regulated Cort response, as monitored by the normal feedback loop through stress-induced Cort and the decrease of *Nr3c1* encoding GR. These mice also displayed no spatial memory impairment, no learning and FC deficits, and no anxiety- or depression-like behaviors. Interestingly, however, they are well protected from PTSD-like memory formation, suggesting therefore that PAI-1 inhibition could be a therapeutic approach for the treatment of this disorder.

### Limitations and strengths

Our study has two major limitations, but also strengths, due to the complementary nature of our translational approach and experimental model. First, our preclinical model explores one dimension of PTSD, namely paradoxical memory impairment. Nevertheless, this dimension is characteristic of this multi-symptomatic pathology, since it is precisely this paradoxical memory characterized by both emotional hypermnesia and contextual amnesia that enables pathological memory to be distinguished from normal fear memory, and which therefore closely mirrors the dissociation observed in PTSD memory processing in humans^11^. By combining contextual FC with a post-conditioning injection of Cort, our model allows us to explore the paradoxical alteration of memory, a cardinal feature of PTSD and listed under the category of “cognitive and avoidance symptoms”^1^. Other symptoms, such as ‘Mood’, ‘Re-experincing’ and ‘hyperarousal and reactivity’, are not assessed by this model. However, treating these memory alterations through the recontextualization of traumatic memory, which has already been shown to suppress traumatic hypermnesia in mice^30^, may also alleviate other symptoms, such as hyperarousal and high anxiety. Interestingly, our human studies showed that blood PAI-1 was correlated with a wider range of PTSD-related psychological distress symptoms, such as perceived stress, anxiety, depression and suicidal thoughts that notably characterize PTSD in humans^87^.

The second limitation of our study concerns the exclusive inclusion of male human participants in the longitudinal study. The reason for this exclusively male composition is due to their belonging to a Special Forces regiment mainly composed of men. The lack of female in the human data analyzed is likely to reduce the scope of our conclusions by potentially underestimating them. Indeed, a meta-analysis of data from the literature indicates that the prevalence of PTSD is significantly higher in women; around twice as high as in men^88^. Interestingly, a comparative study revealed notably that women have significantly higher plasma PAI-1 levels than men, even after adjusting both for age, body mass index and metabolic factors^86^, which could potentially explain their higher prevalence. The strength of the longitudinal study lies in its ability to highlight the evolution of this biological marker by tracking PAI-1 levels in the blood of the same soldiers, both before and after the mission associated with trauma. This approach enables to examine individual differences in symptom trajectories, rather than relying solely on group averages, thereby providing a better understanding of how PTSD evolves in different individuals. A second strength of this study was to be able to correlate PAI-1 with a wide range of PTSD-related symptoms of psychological distress, which characterize PTSD in humans, further strengthening the preclinical blood and hippocampal data and the face validity of the mouse model.

## Conclusion

In conclusion, using both a validated preclinical mouse model of PTSD-like memory and a longitudinal clinical study of human subjects with PTSD symptoms, our data highlight that an elevated blood level of PAI-1 following a traumatic event emerges as a promising biomarker for PTSD, as well as a brain risk factor to develop this pathology. The consistency of these findings across species underscores the potential utility of PAI-1 as a diagnostic tool for PTSD. This biomarker not only reflects the neurobiological changes underlying PTSD but also provides a quantifiable measure of the disorder’s psychological impact, offering new avenues for early intervention and personalized treatment strategies. Furthermore, pharmacological inhibition of PAI-1 could provide an innovative therapeutic avenue for managing this disabling pathology.

## Materials and methods

### Chemicals

For all the experiments we used a preformed water-soluble complex of corticosterone and 2-hydroxypropyl-β-cyclodextrin (#C174, Sigma, USA). In mice, corticosterone (Cort; 2.5 mg/kg) and the vehicle (NaCl 0.9%) were administered intraperitoneally (i.p.) at a volume of 0.1 ml/10 g body weight immediately after the acquisition of fear conditioning in order to mimic the effects of intense trauma (i.e., an hyper-reactivity of the HPA axis)^27^.

### Animals

PAI-1^+/+^ and PAI-1^-/-^ male and female mice were initially obtained from the Jackson laboratory (B6.129S2-Serpine1tm1Mlg/J, Strain #002507, ME USA) and bred in our animal facility. C57Bl/6N male and female mice from Janvier (France) were used for experiments performed only on WT animals. Mice were maintained on a 12 h light/dark cycle (lights on at 8 am/ off at 8 pm) and group-housed in transparent cages with a grid-top with free access to water and food. Three- to four-month-old mice were housed individually one week before experiments began. All experiments were performed in male and female mice according to the protocols approved by the Aquitaine-Poitou Charentes local ethical committee (authorization number APAFlS# 26060-2020061515184837 v3, #30501-2021031816309623_v2 and #51890-2024110516368782_v4) in strict compliance with the French Ministry of Agriculture and Fisheries and European Communities Council Directive (2010/63/EU). All experiments were conducted and analyzed by an experimenter blind to genotype and/or drug treatment conditions; animals were randomly assigned to the experimental groups.

### Restraint Stress

Each mouse was placed into 50 ml conical tube fitted with punctures at its end to allow ventilation. The tubes were placed horizontally, in a clean cage with bedding, under strong light exposure (320 lux) for a continuous period of restraint. After 180 min of restraint, the stressed mice and the unstressed control group (home-cage) were sacrificed by dislocation and their blood and brains were collected.

### Behavioral tests

#### Circadian activity measurements

Mice were placed in actimetry cages (22 x 12 x 18 cm, L x W x H; Imetronic, France) at the end of the day (6 pm) and their activity was recorded for three consecutive days. Mice had access to water and food *ad libitum* with a light/dark cycle similar than their animal facility room. The first day was counted as habituation and was not analyzed. Activity measurements (infrared beams breaks) were acquired from the Imetronic actimetry software (France) and recorded as arbitrary values per hour.

#### Activity in the open field (OF, 30 minutes test)

Mice were placed in an open field (40 x 40 x 40 cm, L x W x H; 120 lux), which was cleaned with 20% ethanol between each trial to remove scents from other animals. Mice were free to explore the open field for 30 minutes and then placed back into their homecage. Ethovision 12 videotracking software (Noldus, Wageningen, Netherland) was used to record the distance traveled during the trial by 5 min blocks.

#### Y-maze (alternation)

Mice were placed in a Y shaped maze (40 x 8 x 12 cm, L x W x H) with a dim light setting (80 lux). The maze was cleaned with 20% ethanol between each trial to remove scents from other animals. Mice were placed in the center of the maze facing a corner and were allowed to explore freely the maze during 8 minutes. Alternance in the maze was scored manually on video and EthoVision 12 videotracking software (Noldus, Netherland) was used to record distance traveled during the test. Percentage of alternation was calculated based on the total number of entries and alternation made: Alternation x 100 / Alternation possibilities (total entries - 2).

#### Y-maze (spatial memory)

Mice were placed in a Y shaped maze (40 x 8 x 12 cm, L x W x H) with a dim light setting (80 lux). The maze was cleaned with 20% ethanol between each trial to remove scents from other animals. Cues on the walls of the testing room provided spatial indications.. One arm was closed with a sliding door and mice were placed in the center of the maze facing a corner and were allowed to explore during 8 minutes. One hour later mice were placed in the same Y-shaped maze but with all three arms open and were allowed to explore during 8 minutes. Exploration was scored manually on video by measuring number of entries in each arm: departure arm (DA), familiar arm (FA) and novel arm (NA) and EthoVision 12 videotracking software (Noldus, Netherland) was used to record distance traveled during the test. Percentage of exploration of NA was calculated by comparing entries in FA and in NA: Entries NA x 100 / (Entries NA + Entries FA).

#### Novel objects recognition (NOR)

Mice were placed in an open field maze (40 x 40 x 40 cm, L x W x H) with a dim light setting (25 lux). The maze was cleaned with 20% ethanol between each trial to remove scents from other animals. The first day, mice were free to explore the open field for 15 minutes and then placed back into their homecage (habituation). On the second day two identical objects were placed in the open field and mice were allowed to explore for 10 minutes (acquisition). The third day, one of the two objects was changed to a novel object (NO) different from the familiar object (FO) in shape and color (memory test). Mice were allowed to explore for 10 minutes these two objects and EthoVision 12 videotracking software (Noldus, Netherland) was used to record distance traveled and time spent exploring each object. Percentage of NO exploration was calculated using time exploring the NO over total exploration time: Time NO x 100 / (Time NO + Time FO).

#### Cued fear conditioning (FC) and extinction

Mice were pre-exposed to the context A consisting of a white PVC chamber (30 x 24 x 22 cm, L x W x H) with an opaque PVC floor, for 2 min, in a brightness of 10 lux with a scent of acid acetic 1%. The next day acquisition of fear conditioning was performed in context B, consisting in a transparent conditioning chamber (30 x 24 x 22 cm, L x W x H) with the floor connected to a shock generator, in a brightness of 110 lux allowing for mice to see visual cues present in the testing room, with a scent of ethanol 70%. Briefly, each animal placed in the conditioning chamber for 4 min received 2 footshocks (0.4 mA, 50 Hz, 1 s) co-occurring with the end of the two tones deliveries (70 dB, 1 kHz, 15 s). After fear conditioning, each animal was returned to its home cage and the next day, all mice were submitted to the tone memory tests during which freezing behavior, defined as a lack of any movement except for respiratory-related movements, was measured and used as an index of conditioned fear. Mice were re-exposed to the tone in the context A during which three successive recording sessions of the behavioral responses were performed: one before (first 2 min), one during (next 2 min), and one after (last 2 min) tone presentation. The next 3 following days mice were placed back in the context A for the same tone memory test to measure extinction of acquired fear memory. Animals freezing was recorded and scored using automatic counting software (Imetronic, France). Freezing values represented in Tone and Ext 1, 2 and 3 (**Fig. 2D, left panel**) are freezing levels during sound presentation only.

#### Contextual fear conditioning (FC) and extinction

Acquisition of fear conditioning was performed in a transparent conditioning chamber (30 x 24 x 22 cm, L x W x H) with the floor connected to a shock generator, in a brightness of 110 lux allowing for mice to see visual cues present in the testing room, with a scent of ethanol 70%. Briefly, each animal was placed in the conditioning chamber for 4 min and received 2 footshocks (0.4 mA, 50 Hz, 1 s). After fear conditioning, each animal was returned to its home cage and the next day, all mice were submitted to the context memory test during which freezing behavior, defined as a lack of any movement except for respiratory-related movements, was measured and used as an index of conditioned fear response. On the next three days, mice were placed back in the conditioning context during 6 minutes for a context memory test, first to evaluate acquisition of the contextual conditioning and then the extinction of acquired fear memory. Animals freezing was recorded and scored using automatic counting software (Imetronic, France). Freezing values represented in Ext 1, 2 and 3 **(Fig. 2D, right panel)** are freezing levels during the 6 minutes of context re-exposure test.

#### Stress-induced hyperthermia (SIH)

SIH was assessed by measuring the effect of an acute stress on body temperature. Acute stress was the rectal temperature measurement which was done using a lubricated rectal thermometer probe. Temperature was measured once a day for four days at the same time to ensure that the measurement on the test day would not be affected by any environmental factor. This also served as habituation to the manipulation. On the fourth day (test day), another temperature measurement was made after 10 minutes to assess the hyperthermia induced by a stressor. Delta between T10 and T0 measurements represents SIH (T10-T0).

#### Elevated-plus maze (EPM)

The EPM assay is a widely used behavioral test to measure anxiety-like behavior in rodents. Mice were placed in an elevated (60 cm) plus maze composed of two open (OA) and two closed (CA) arms (35 x 6 x 15 cm, L x W x H) with a dim light setting (50 lux) and was cleaned with 20% ethanol between each trial to remove scents from other animals. Mice were placed in the center of the maze, head facing an open arm and were free to explore the EPM for 5 minutes and then placed back into their home cage. EthoVision 12 videotracking software (Noldus, Netherland) was used to record the time spent in each zone during the trial. Mice were considered to be in a zone when the four paws were inside. Three zones were defined using the tracking software: one in the center of the maze, two in the open arms (anxiogenic space) and two in the closed arms. Anxiety-like behavior is expressed as the percentage of time spent in the anxious area (OA): (time in OA / time in OA + time in CA) x 100.

#### Open field (OF)

The OF assay quantifies anxiety-like behavior primarily based on rodents’ avoidance of the open center and preference for the protected periphery. Mice were placed in an open field (40 x 40 x 40 cm, L x W x H, 120 lux) cleaned with 20% ethanol between each trial to remove scents from other animals. Mice were free to explore the open field for 5 minutes and then placed back into their home cage. EthoVision 12 videotracking software (Noldus, Netherland) was used to record distance traveled and time in zone during the trial. Mice were considered to be in a zone when the four paws were inside. Two zones were defined using the tracking software: a central zone (20 x 20 x 20 cm, L x W x H anxiogenic space) and a peripheral zone (10 cm wide zone, protected space).

#### Light/dark box (LD)

The LD assay is also a classical behavioral assay to measure anxiety-like behavior in rodents. Mice were placed in a rectangular box (38 x 19 x 15 cm, L x W x H) containing one light area (19 x 19 x 15 cm, L x W x H, 340 lux) and one dark area (19 x 19 x 15 cm, L x W x H, 3-5 lux) separated by a wall (15 cm, H) with an open door allowing it to go from one area to the other. The maze was cleaned with 20% ethanol between each trial to remove scents from other animals. Mice were placed in the light area head facing a wall and were free to explore the LD box for 5 minutes and then placed back into their home cage. EthoVision 12 videotracking software (Noldus, Netherland) was used to record time in the bright light area during the trial as an index of anxiety due to mice’s innate aversion to bright environment. Mice were considered to be in a zone when the four paws were inside. Two zones were defined using the tracking software: light and dark areas.

#### Forced swim test (FST)

Mice were placed in a transparent Plexiglass cylinder (Ø12.5 cm, H24.5 cm) containing 24°C water with a dim light setting (90 lux) for 6 minutes. At the end of a trial, mice were placed in a cage to dry under a warm light and then returned to their home cage. Water was cleaned between each trial and warm water was added to keep temperature at 24°C. Time spent immobile, as an index of “behavioral despair” reflecting depressive-like states, was scored manually. Only the four last minutes of the test were used (the first two minutes were designated as habituation).

#### Sucrose preference (SP)

The SP test is a measure of anhedonia, which is the reduced ability to experience pleasure and is a core symptom of depressive-like states. SP was performed directly in the home cage where two bottles were placed on the top of it: one containing water and the other containing a 2% saccharose (wt/vol) solution in water. Each day for 5 days, both bottles were weighed to measure consumption, and their place was reversed each day to avoid any place preference effect. SP was measured during the last 4 days of measurement as the first 24h was considered as habituation. SP (%) was calculated using this formula: (saccharose consumption / total liquid consumption) × 100.

#### Adaptive vs maladaptive (PTSD-like) fear memory using contextual fear conditioning

The behavioral model based on a general fear conditioning procedure has been fully described in previous studies^18,27,30,89,90^. It is composed of 3 main phases:

**1) Pre-exposure**: the day before fear conditioning, each mouse was placed individually in a transparent PVC chamber (30 x 40 x 32 cm, L x W x H) with an opaque PVC floor, for 2 min, in a brightness of 10 lux with a scent of acid acetic 1%. This pre-exposure allowed the mice to acclimate and become familiar with the chamber used for the cue alone test (“safe context”).
**2) Induction of adaptive vs PTSD-like fear memory**: acquisition of fear conditioning was performed in a different context, consisting of a transparent conditioning chamber (30 x 40 x 32 cm, L x W x H) containing visual cues on each wall and with the floor connected to a shock generator, under a brightness of 110 lux with a scent of ethanol 70%. Briefly, each animal was placed in the conditioning chamber for 4 min and received 2 footshocks (0.4 mA, 50 Hz, 1 s), which never co-occurred with two tone deliveries (70 dB, 1 kHz, 15 s). This tone-shock unpairing paradigm is known to make contextual cues the primary stimuli associated with the footshock ^27^. Consequently, the phasic tone, although salient, is not predictive of the shock delivery, whereas the static contextual cues constitute the main predictor of the shock. Immediately after the acquisition of fear conditioning, mice received a systemic i.p. injection of either NaCl or Cort (2.5 mg/kg). An adaptive fear memory is therefore attested in control mice (NaCl-injected) by the expression of highly conditioned fear when re-exposed to the conditioning context, as well as their lack of conditioned fear when re-exposed to the irrelevant tone cue (in the safe context). In contrast, Cort-injected mice display a maladaptive (PTSD-like) memory attested by an abnormally high fear response to the irrelevant tone (cue-based hypermnesia) together with a decreased conditioned fear to the conditioning context (contextual amnesia) ^27^.
**3) Memory tests**: after fear conditioning, each animal was returned to its home cage and 4-5 days later, all mice were submitted to two memory tests during which freezing behavior, defined as a lack of any movement except for respiratory-related movements, was measured and used as an index of conditioned fear. During these two memory tests, animals were continuously recorded for off-line second-by-second scoring of freezing by an observer blind to the experimental groups. Mice were first re-exposed to the tone within the “safe” context during which three successive recording sessions of the behavioral responses were performed: one before (first 2 min), one during (next 2 min), and one after (last 2 min) tone presentation. Conditioned response to the tone is expressed by the percentage of freezing during the tone presentation compared to the levels of freezing expressed before and after tone presentation (repeated measures on 3 blocks of freezing). The strength and specificity of this conditioned fear are assessed by a ratio that represents the increase in the percentage of freezing during the tone relative to a baseline freezing levels {i.e., pre- and post-tone periods mean: (tone - ((pre + post)/2)) / (tone + ((pre + post)/2))}. Then 24h later, mice were re-exposed to the conditioning context alone for 6 min (without the tone cue). Freezing to the context was calculated as the percentage of the total time spent freezing during the successive three blocks of 2-min periods of the test. While the first block is the critical block attesting difference between animals during retrieval, the following two blocks are presented in order to assess a gradual extinction of the fear responses in the absence of shock.

A PTSD score was calculated for each experiment using this model. This score was the mean of two z-scores: one based on the tone fear response (tone ratio = x-tone) and the other based on the context fear response (100-Cx 0-2 = x-context). To calculate z-score this formula was used based on a published article for behavioral representation^91^: ((x-tone or x-context) – (mean x-tone or x-context of the NaCl groupe)) / (standard error of the mean or sem x-tone or x-context of the NaCl groupe).

### Quantitative PCR analysis

Samples from different brain structures (dorso/ventral hippocampus, amygdala and medial prefrontal cortex) were homogenized in Tri-reagent (Euromedex, France) and RNA was isolated using a standard chloroform/isopropanol protocol. RNA was processed and analyzed using an adapted version of published methods^92^. cDNA was synthesized from 2 μg of total RNA using RevertAid Premium Reverse Transcriptase (Fermentas, Thermo Fisher Scientific, USA) and primed with oligo-dT primers (Fermentas, Thermo Fisher Scientific, USA) and random primers (Fermentas, Thermo Fisher Scientific, USA). qPCR was performed using a LightCycler® 480 Real-Time PCR System (Roche, Meylan, France). qPCR reactions were done in duplicate for each sample, using transcript-specific primers, cDNA (4 ng) and LightCycler 480 SYBR Green I Master (Roche) in a final volume of 10 μl. The PCR data were exported and analyzed in a computer-based tool (Gene Expression Analysis Software Environment) developed at the Neurocentre Magendie (France). The Genorm method was used to determine the reference gene. Relative expression analysis was corrected for PCR efficiency and normalized against two or three reference genes. *Gapdh*, *Eef1a1*, *Ppia*, *Atp5f1b* and *Ube2d2a* were used as reference genes in the different qPCR experiments. The relative level of expression was calculated using the comparative (2-ΔΔCT) method and then represented as fold change from the control groups (i.e., non-stressed and NaCl-injected groups). qPCR amplification used specific primers to specifically amplify *Serpine1* encoding PAI-1 protein, *Nr3c1* encoding glucocorticoid receptor (GR), *Gapdh*, *Eef1a1*, *Ppia*, *Atp5f1b* and *Ube2d2a* as reference genes.

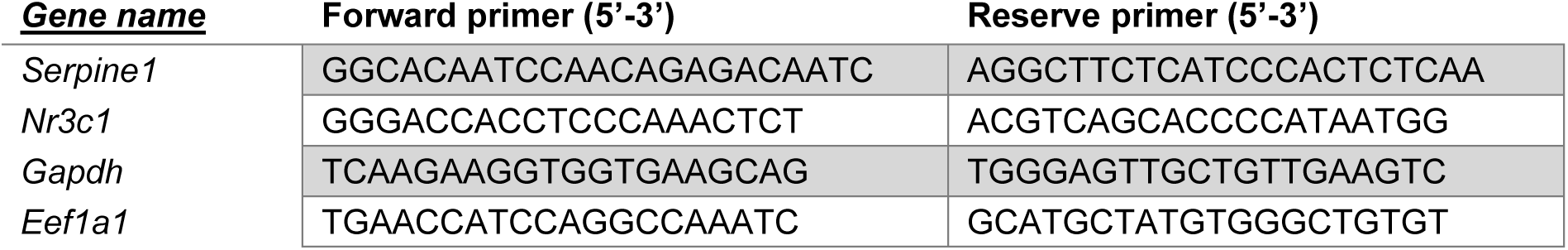
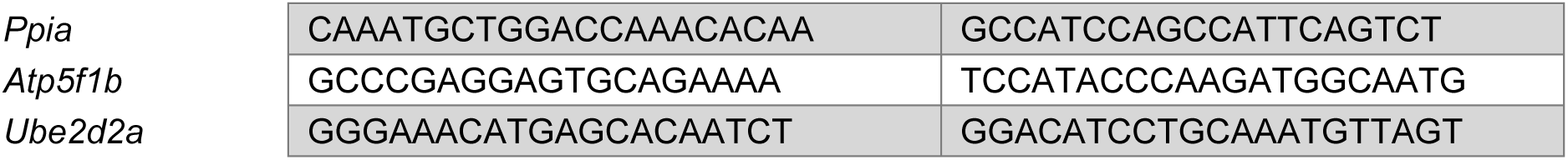

### Single-molecule fluorescence *in situ* hybridization (RNAscope)

For the examination of targeted RNA within intact cells, *in situ* hybridization RNAscope technology was used following the protocol described by the supplier. Brain tissue was frozen immediately on dry ice and stored at −80 °C. Brains were sectioned at −17 °C using a cryostat at a thickness of 16 μm and mounted onto Superfrost Ultra Plus slides (Thermo Scientific, J4800AMNZ). Brain sections were collected for dorsal hippocampus from bregma −1.35 mm and −2.2 mm. Probes for *Serpine1* (ACDBio, 402501-C1) was used with the RNAscope Fluorescent Multiplex Kit (ACDBio; 320850) as described by the supplier. Slides were counterstained for DAPI and mounted with ProLong Diamond Antifade mountant (Invitrogen, P36961). Images covering dorsal hippocampus and images from each region of interest were acquired using sequential laser scanning confocal microscopy (Leica SP8). Histograms show *Serpine1*-positive spot density (dots/mm²), quantified from 3–4 bilateral dorsal hippocampal images per mouse (n=4 mice). Regions of interest (ROIs) were delineated in Fiji ImageJ, and *Serpine1*-positive dots within ROIs were counted using the Find Maxima plugin.

### Human soldier cohort Population

The study was conducted on a population of 80 male soldiers who were deployed for six months in Afghanistan in the spring of 2011. The subjects were volunteer for participating to the study, aged between 18 and 50 years and medically fit for military deployment. The study was conducted in accordance with all applicable regulatory requirements. It was approved by CPP SOUTHEAST V Ethics Committee in France, which is registered with the French Department of Health and Human Services (identification number 2010-A01232-37). All volunteers provided written informed consent before participation.

### Protocol

The objectives of the investigations were explained by the military health authorities during a briefing carried out approximately 1 month before a 6-month war-zone deployment. Data were obtained twice: once between February and March 2011, during preparations for the deployment (first run of blood collection) and second 6 months after the end of the mission within the regiment (second run of blood collection). At inclusion, subjects participating to the investigation completed a 1-hour set of “paper and pencil” standardized assessments including common socio-demographic data, psychological and pathological functioning. They were asked to come on the following morning to the medical center for blood collection.

### Psychological variables

#### Socio-demographic evaluation

The socio-demographic variables included age, gender, marital status, tobacco use and experience as a military soldier in duration of duty, antecedent of overseas deployment and, when positive, the number of deployments.

#### Psychological evaluation

The **Freiburg Mindfulness Inventory (FMI)**^93,94^ was used to assess mindfulness (14 items, four-point Likert scale ranging from 1 = strongly disagree to 4 = strongly agree). This questionnaire involves both dimensions (i.e., presence and acceptance) and is applicable to all population groups, in particular those who do not practice mindfulness meditation. The higher the score is, the more mindful the participant is.

The **Siebold vertical and horizontal cohesion** questionnaire assesses cohesion at the platoon level using four items for horizontal bonding among peers and four items for vertical bonding between leaders and subordinates^95^. Higher scores indicate greater group cohesion. The **Positive Affect and Negative Affect Scale (PANAS)**^96^ is a good index for distress evaluation when positive affects (PA) are below 33.3 and negative affects (NA) above 17.4^97,98^.

#### Stress and mental health evaluation

Although the subjects were declared healthy after a medical examination, they fulfilled the Post-traumatic Stress Disorder Check List, the Cohen perceived stress, the Hospital Anxiety-Depression Scale and the General Health Questionnaire to assess how they were adapted to the environment and scheduled mission.

The **PTSD CheckList (PCL-S)** was used to detect post-traumatic stress disorder (PTSD) according to DSM-IVR (American Psychiatric Association, 1994). The usual cut-off for PCL-S is at ≥ 44 ^36^. However, for French military personal, a cut-off of 34 is considered as sensitive and specific for this specific population^99^.

The **Cohen Perceived Stress** was evaluated using the Perceived stress scale (PSS)^100^ with a French validated version^101,102^. PSS is a self-reported measure which assesses the degree to which the respondent has perceived situations in his/her life within the past month as stressful. Higher values indicating greater perceived stress.

The **Hospital Anxiety and Depression Scale (HADS)** aimed to detect anxiety and depression in general non-psychiatric medical outpatients^103,104^.

The mental suffering was checked with the **General Health Questionnaire (GHQ-28)** in the 28 items form usually used in healthy population for epidemiological description^105^, as its French validated version^106^. A score over 22 highlights a psychological distress^107^. A four-item suicidal ideation subscale of the GHQ-28 has been used to assess suicidal ideation^108^.

For all questionnaires, data were excluded when more than two items were not completed. When only one item was not completed, it was affected by the value of the mean of the other items.

### Blood collection and ELISA assays

#### Mice blood collection and ELISA

Blood was rapidly collected from the submandibular vein in heparin-EDTA-coated tubes (Sarstedt, France) and centrifuged at 10,000 rpm (4°C, 10 min). Supernatant containing the blood plasma was stored at −80°C, and then processed for ELISA assay. Plasma PAI-1 (Mouse PAI-1 Total Antigen ELISA KIT #MPAIK-TOT, Innovative Research, MI, USA) and corticosterone (Corticosterone EIA kit #K014-H1, Arbor Assays, MI, USA) levels were quantified by ELISA following the manufacturer’s instructions.

#### Human blood collection and ELISA

In the human cohort, blood was collected on dry tube at inclusion (i.e., before) and 6 months after returning from a 6-month mission in a war zone (i.e., after). For each session of blood collection, subjects were asked to not practice sport, drink coffee or smoke 2 hours before blood collection. The morning blood collection was practiced between 8h30 and 10h30 to control circadian variations, and after ten minutes spent in a calm area. Serum PAI-1 (Human PAI-1 Total Antigen ELISA KIT #HPAIK-TOT, Innovative Research, MI, USA) levels were quantified by ELISA following the manufacturer’s instructions.

### Data analysis

#### Mice data analysis

In this study we used both males and females and data represent both sexes pooled together. For behavioral studies, some results were expressed as a percentage of control in order to normalize baseline differences and allow data from male and female groups to be pooled, despite behavioral differences associated with gender (i.e., actimetry, open field activity, forced swim test and sucrose preference). Sample size (n=6-30) were chosen to ensure adequate statistical power for all experiments. Dependent variables were checked for normal distribution (Shapiro-Wilk test, *p>0.05*). Statistical analyses of normally distributed variables were performed using Student’s t-tests were used for pairwise comparisons and one-way or two-way analyses of variance (ANOVA) were used (depending on the number of independent variables) when comparing more than two groups. Repeated measurements analysis was used if more than one measure originated from the same animal. These ANOVA were followed by Fisher’s PLSD post-hoc test for pairwise comparisons when necessary. Non normally distributed variables were analyzed using non parametrical tests: a Mann-Whitney test for pairwise comparison and a Kruskal-Wallis test for one-way ANOVA. A significance level of *p<0.05* was used for all statistical analyses. Statistical significance was expressed as \**p<0.05*; \*\**p<0.01*; \*\*\**p<0.001* and ^##^ *p<0.01*; ^###^ *p<0.001* for genotype effect and differences from chance. All data (bar graphs or plots) were expressed as mean ± s.e.m. Correlation p-values were calculated with either Pearson (normally-distributed, parametric) or Spearman (not normally distributed, non-parametric) coefficients, both of which are marked as r^2^. All analyses were performed using GraphPad Prism 7 software (San Diego, CA, USA).

#### Human data analysis

In this study, 80 male soldiers were included from whom blood samples were taken 1 month before a 6-month mission in a war zone and 6 months after their return from the mission. At inclusion, they answered to several questionnaires (i.e., mindfulness, cohesion, perceived-stress, hospital anxiety and depression, general health questionnaire) including PCL-S, which is the standardized method to assess PTSD symptoms in the military. To distinguish soldiers with PTSD symptoms from control soldiers with low level of PTSD symptoms a cut off at a PCL-S score of 34 was used. It was previously described in military cohorts^99^ and present a 78% sensitivity and 94% specificity to detect subjects in need of psychotherapy and/or pharmacological treatment related to traumatic experiences. To measure the evolution of blood PAI-1 over a one-year period, we calculated **ΔPAI-1 = PA-1 level after – PAI-1 level before**. To evaluate the potential of PAI-1 as a diagnostic tool we quantified true positive (TP), false positive (FP), true negative (TN) and false negative (FN) using different cut off values and used formula for sensitivity (Se = TP / [TP + FN]), specificity (Sp = TN / [TN + FP]), negative (NPV = TN / [TN + FN]) and positive predictive values (PPV = TP / [TP + FP]), and Youden index (J = Se + Sp – 1). Youden index gives an estimated value between 0 and 1 or 0 and −1 representing a biomarker validity by taking into account both specificity and sensitivity^37^. A significance level of *p<0.05* was used for all statistical analyses. Statistical significance was expressed as \**p<0.05*; \*\**p<0.01*; \*\*\**p<0.001* and bar graph expressed as mean ± sem. Correlation p-values were calculated with either Pearson (normally-distributed, parametric) or Spearman (not normally distributed, non-parametric) coefficients, both marked as (r). All analysis were performed with Graph Pad Prism 7 software (San Diego, CA USA).

## Acknowledgments

We thank all the members of the PUMA platforms of the Magendie Neurocentre in particular Delphine Gonzales, Elisabeth Huc, Sara Laumond and Julie Tessaire of the genotyping and animal’s facility for mouse breeding and care; Thierry Lesté-Lasserre from the transcriptomics platform for qPCR analyses. We also thank Yann Rufin from the Biochemistry and Biophysics (BioProt) Facility of the Bordeaux Neurocampus (LABEX BRAIN; ANR-10-LABX-43). We also thank Dr Muriel Koehl and Dr Lyès Kaci for developing the PTSD score. This work was financially supported by Inserm (3U1215E011SEREVEST), Inserm Transfert (CoPoC 1 #MAT-PI19520-A-01 and CoPoC 2 #MAT-PI19520-A-04), Fondation de France (00147198/WB-2023-49762), University of Bordeaux and Agence Nationale pour la Recherche via the SPARK program (ANR-21-EXES-0004). The funders had no role in study design and data analysis and nor in the decision to submit the work for publication.

## Author contributions

JMR conceived the project; JMR and MT supervised the project; MM and JMR designed the experiments; MM, SA, VRL, ST, AC, VL, PRL, LM, EV performed the experiments; MM, EV, AMD, DC, MV, AD, MT and JMR analyzed and interpreted the data; MM and JMR wrote the manuscript. All authors approved the manuscript and declare no competing interests.

---

**Supp. Fig. 1.**
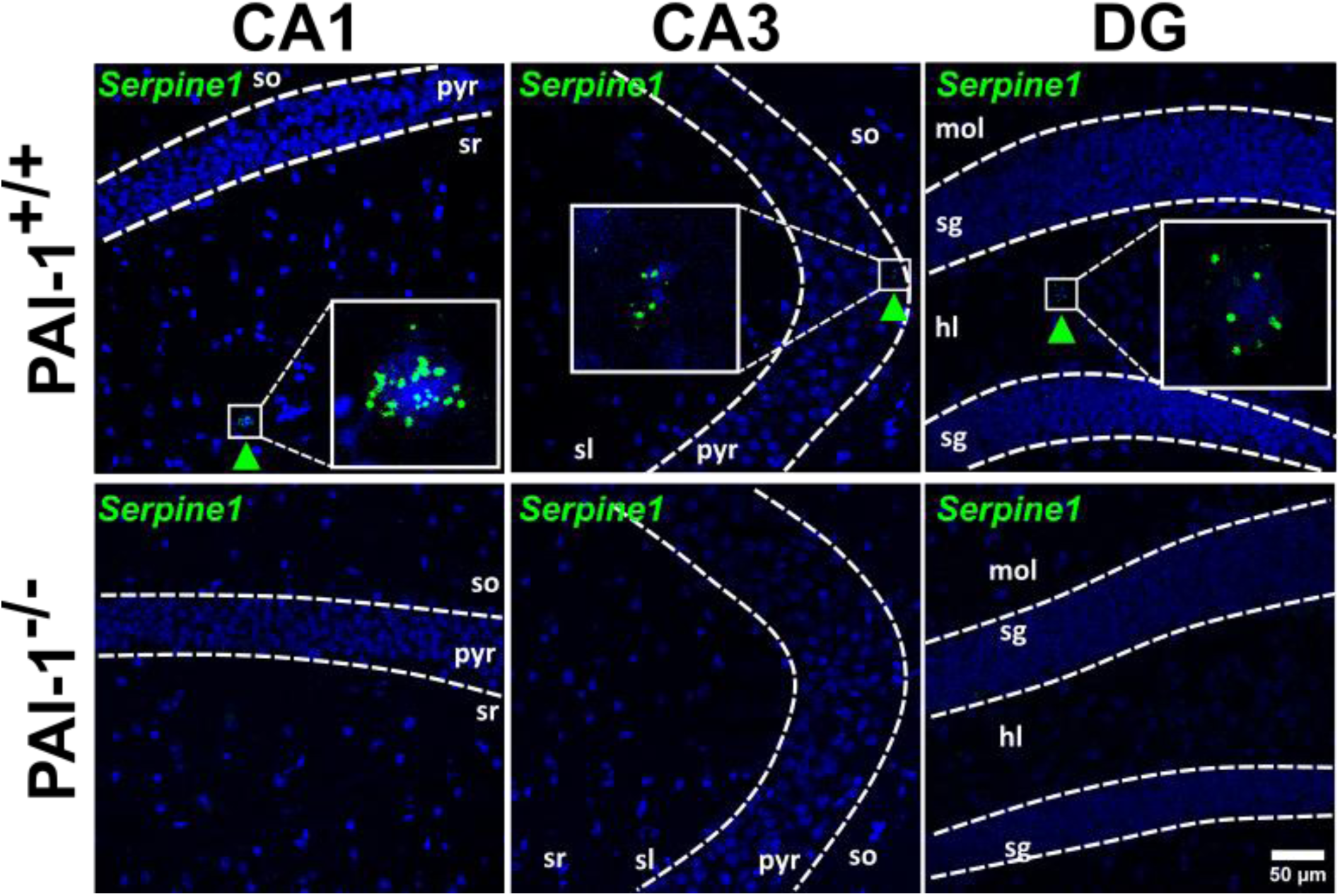
PAI-1-deficient (PAI-1^-/-^) male and female mice do not express PAI-1 in the hippocampus. Confocal images of coronal brain sections of the dorsal hippocampus (Hpc) of PAI-1^+/+^ and PAI-1^-/-^ mice showing the distribution of *Serpine1* encoding PAI-1 (green) detected by single-molecular fluorescent *in situ* hybridization (RNAscope, top panel). No PAI-1 expression is detected in the Hpc of PAI-1^-/-^ mice (bottom panel). Slides were counterstained with DAPI (blue). <u>Pyr</u>: pyramidal cell layer; <u>so</u>: stratum oriens; <u>sr</u>: stratum radiatum; <u>sl</u>: stratum lucidum; <u>mol</u>: molecular layer of the DG; <u>sg</u>: stratum granulosum; <u>hl</u>: hilus of the DG.

